# Severity of Addiction-like Behaviors and Sex Distinctly Promote Oxycodone-Related Behaviors in Heterogeneous Stock Rats

**DOI:** 10.64898/2026.11.24.768451

**Authors:** Sara Foroutani, Elizabeth A. Sneddon, Joseph Mosquera, Alex Morgan, Jarryd Ramborger, Benjamin C. Sichel, Dyar N. Othman, Abraham A. Palmer, Leah C. Solberg Woods, Giordano de Guglielmo, Lieselot L.G. Carrette, Marsida Kallupi, Olivier George

**Affiliations:** Department of Psychiatry, University of California, San Diego, La Jolla CA 92093, USA; Institute for Genomic Medicine, University of California San Diego, La Jolla CA 92093, USA; Department of Internal Medicine, Molecular Medicine, Center on Diabetes, Obesity and Metabolism, Wake Forest School of Medicine, Winston-Salem, NC, USA

**Keywords:** sex differences, individual differences, Heterogeneous Stock rats, estrous cycle, oxycodone, opioids, addiction-like behavior

## Abstract

The opioid epidemic continues to impose substantial public health burdens. Critical gaps remain in understanding how sex and individual differences contribute to opioid addiction-like behavior. Although previous clinical and preclinical work has identified sex and other individual differences in opioid-related outcomes, it remains unclear whether the magnitude of sex differences varies across behavioral domains and levels of individual addiction-like vulnerability.

To address this gap, we analyzed data from 810 genetically diverse heterogeneous stock (HS) rats that underwent oxycodone self-administration under short-access (2h/day) and extended-access (12h/day) conditions. Addiction-like behavior was characterized across four domains comprising escalation of oxycodone intake, motivation to obtain oxycodone, tolerance to oxycodone-induced analgesia, and withdrawal-induced hyperalgesia. These measures were integrated into a composite Addiction Index (AI) to characterize individual addiction-like severity, with animals classified into four groups: resilient, mild, moderate, and severe. Sex differences were then examined across behavioral outcomes, AI groups, and experimental cohorts. Estrous-phase effects on motivation, relationships among behavioral dimensions, and within-session patterns of oxycodone intake were also examined.

Females exhibited greater oxycodone intake and motivation than males following extended access, whereas pain-related behavioral outcomes showed greater associations across levels of addiction-like severity than between sexes. Although sex differences were observed for specific behavioral outcomes, further analyses provided no evidence that their magnitude varied across AI groups. Estrous phase did not significantly affect motivation for oxycodone. Pooled sex effects supported greater self-administration and motivation in females, whereas pooled sex effects for pain-related outcomes were not significant and showed varying degrees of between-cohort heterogeneity. Temporal features of oxycodone intake also differed by sex and were associated with addiction-like severity, with titration-phase intake showing the strongest association with the AI. Relationships among addiction-related behavioral dimensions were largely similar between females and males, although specific associations differed by sex and across AI groups.

These findings demonstrate that sex and individual variability contribute in different ways to oxycodone addiction-like behavior. Female-biased differences were most apparent following extended access, whereas pain-related behavioral outcomes showed greater associations across levels of addiction-like severity. Together, these results highlight the importance of multidimensional phenotyping for characterizing sex and individual variability in opioid addiction-like behavior.

## Introduction

The opioid epidemic remains a critical public health crisis in the United States, with more than 80,000 opioid-involved overdose deaths reported annually and more than 6.1 million people meeting diagnostic criteria for opioid use disorder (OUD) in 2022 (SAMHSA, 2023; Dowell et al., 2024). Although men historically exhibit higher rates of opioid misuse and overdose (Butelman et al., 2023), women show a more rapid progression from prescription opioid misuse to dependence and greater craving (Back et al., 2011; Becker & Koob, 2016; McHugh et al., 2013). Together, these findings underscore the importance of understanding how sex contributes to variability in opioid-related behaviors.

Sex differences in opioid responses may also be influenced by ovarian hormonal state. Clinical studies have primarily examined menstrual-cycle effects on opioid analgesia rather than opioid use, with mixed findings across drug types and experimental conditions. Some studies have observed increased analgesic effects during the follicular phase, when estrogens rise, (Ribeiro-Dasilva et al., 2011), whereas others have reported no phase-dependent differences (Ahmed et al., 2012; Ribeiro-Dasilva et al., 2011; Sener et al., 2005). Preclinical research also exhibits sex differences across multiple opioid-related behaviors, with females showing a more rapid escalation of opioid self-administration and greater motivation to obtain opioids under several experimental conditions. Estrous-dependent variation in opioid self-administration has also been reported, although findings regarding ovarian hormone effects are mixed. Several studies have observed reduced opioid intake during proestrus or estrus (Hinds et al., 2023; Patel & Loweth, 2024; Sharp et al., 2021), whereas others have reported no change or increased responding across estrous phases or following hormonal manipulation (Cicero et al., 2003; D’Ottavio et al., 2023; Lynch & Carroll, 1999; Nogueira et al., 2024; Roth et al., 2004; Towers et al., 2023). However, the influence of hormonal state on OUD remains largely underexplored in clinical research, despite associations between ovarian hormones and opioid-related behaviors (Maher et al., 2023) and, for preclinical studies, these have primarily used conventional laboratory strains, leaving the role of estrous state less understood in genetically diverse populations. Heterogeneous Stock (HS) rats provide an opportunity to examine sex and estrous-related differences in the context of substantial genetic and behavioral variability. Derived from eight inbred founder strains, HS rats encompass broad genetic and phenotypic variability that more closely reflects the diversity observed in human populations and have therefore been used to investigate individual differences in addiction-related behaviors (Chitre et al., 2020; Hansen & Spuhler, 1984; Solberg Woods & Palmer, 2019). Recent studies have leveraged this population to identify genetic, transcriptional, and circuit-level correlates of addiction-related behavioral variability and to characterize individual differences in addiction-like phenotypes across several drugs, including oxycodone (Allen et al., 2021; Cannella et al., 2024; Carrette et al., 2021; Delorme et al., 2025; Kallupi et al., 2020, 2026; Kuhn et al., 2025a, 2025b; Lara et al., 2026; Sedighim et al., 2021). Oxycodone addiction vulnerability is multidimensional, with escalation of drug intake, motivation, analgesic tolerance, and withdrawal-induced hyperalgesia capturing distinct features in HS rats, that can be integrated into a composite measure of addiction severity (George & Koob, 2017; Kallupi et al., 2020, 2026).

We previously showed that severe addiction-like behavior is in general associated with greater initial analgesia, escalation, and tolerance compared with resilient, with females exhibiting greater escalation and motivation than males but similar tolerance and hyperalgesia. Individually differential relationships among these measures further support heterogeneity in the behavioral profiles underlying overall addiction vulnerability (Kallupi et al., 2026). Thus, population-level sex comparisons may not capture how sex differences are expressed across the spectrum of addiction vulnerability. Whether the magnitude of sex differences varies across addiction-like vulnerability levels or experimental cohorts, and whether relationships among these addiction-related dimensions vary by sex or vulnerability, remains unclear. Building on previous characterization of oxycodone addiction-like behavior in HS rats (Kallupi et al., 2020, 2026), the present study used HS rats to examine how sex and individual variability relate to oxycodone addiction-like behavior, and how reliable sex differences are between independent cohorts. We hypothesized that sex differences would vary across AI-defined resilient, mild, moderate, and severe groups, that motivation would vary across estrous phases in females, and that temporal features of oxycodone intake would be associated with overall addiction-like severity.

## Methods

### Subjects

Outbred Heterogeneous Stock (HS) rats were obtained from Dr. Leah Solberg Woods at Wake Forest University School of Medicine (WFU; NMcwiWFsm #13673907, RRID: RGD_13673907) and Dr. Abraham Palmer at UC San Diego School of Medicine (UCSD; McwiWfsmAap: HS #155269102, RID: RGD_155269102). Male and female HS rats (N = 810; females: n = 387, males: n = 423) were used across 18 experimental cohorts. Upon arrival at 3–4 weeks of age, animals from WFU underwent a two-week quarantine. Following quarantine, rats were pair-housed in a temperature- and humidity-controlled vivarium (20–22 °C, 45–55% humidity) under a reversed 12-h light/dark cycle. Rats from UCSD did not have to undergo a quarantine period. Rats had *ad libitum* access to tap water and standard chow (PJ Noyes Company, Lancaster, NH, USA). All procedures followed the National Institutes of Health Guide for the Care and Use of Laboratory Animals and were approved by the Institutional Animal Care and Use Committee of the University of California, San Diego.

### Drugs

Oxycodone hydrochloride was provided by the National Institute on Drug Abuse (Bethesda, MD, USA) and dissolved in 0.9% sterile saline. Intravenous infusions were delivered at 150 µg/kg per infusion, a regimen previously shown to produce plasma oxycodone concentrations of approximately 40 ng/mL (Kallupi et al., 2020; Kimbrough et al., 2020; Mavrikaki et al., 2017; Wade et al., 2017).

### Intrajugular catheterization

Rats were anesthetized with isoflurane (1–5%) for aseptic surgical placement of catheters in the right jugular vein (de Guglielmo et al., 2024; Kallupi et al., 2020, 2022). Catheters were constructed from Micro-Renathane tubing (18 cm length, 0.023-in inner diameter, 0.037-in outer diameter; Braintree Scientific, Braintree, MA, USA) attached to a 90° angled guide cannula (Plastics One, Roanoke, VA, USA). The catheter was secured to the cannula with dental acrylic and mesh (1 mm thick, 2 cm diameter). Following venous puncture with a 22-gauge needle, tubing was inserted and sutured in place. The guide cannula was routed subcutaneously to an incision on the back and capped to maintain sterility. Preoperative analgesia was administered with flunixin (2.5 mg/kg), and cefazolin (330 mg/kg) was provided as antibiotic during postoperative care. Animals recovered for seven days before behavioral testing. During recovery, catheters were flushed daily with heparinized saline (10 U/mL heparin sodium in 0.9% bacteriostatic saline) containing cefazolin (52.4 mg per 0.2 mL).

### Estrous cycle monitoring

Estrous cycle phase in female rats was assessed at baseline, after the short-access phase during the first progressive ratio session (ShA PR), and after the long-access phase during the second progressive ratio session (LgA PR). Vaginal smears were collected, and light microscopy using a BZ-X800 Analyzer (Keyence, Itasca, IL) was conducted to identify phases based on cytological characteristics (Sneddon et al., 2026).

### Operant self-administration

As previously described in Kallupi et al., 2020, escalation of oxycodone intake was evaluated using an intravenous self-administration paradigm conducted in operant conditioning chambers (29 × 24 × 19.5 cm; Med Associates, St. Albans, VT, USA) housed in sound-attenuating, ventilated cubicles. Chambers contained transparent front doors and back walls, with the remaining walls made of opaque metal. Two retractable levers were mounted on the left side panel and extended at session onset. Presses on the right (active) lever delivered oxycodone (150 µg/kg per infusion in saline) through an intravenous catheter connected to an infusion pump operating under a fixed ratio 1 schedule. Each press resulted in an infusion of 0.1 mL of drug over six seconds, followed by a 20-second timeout signaled by a cue light above the lever; responses during the timeout had no programmed consequences. The left (inactive) lever was recorded but not reinforced. Fluid delivery and behavioral data were recorded using MED-PC IV software.

Rats underwent four daily short-access (ShA) sessions (2 h/day), followed by 14 long-access (LgA) sessions (12 h/day) conducted Monday through Friday during the dark phase. During LgA sessions, animals had *ad libitum* access to food and were provided with a wooden block (3 cm × 3 cm × 3 cm) for enrichment (**Fig. 1A**).

**Figure 1.**
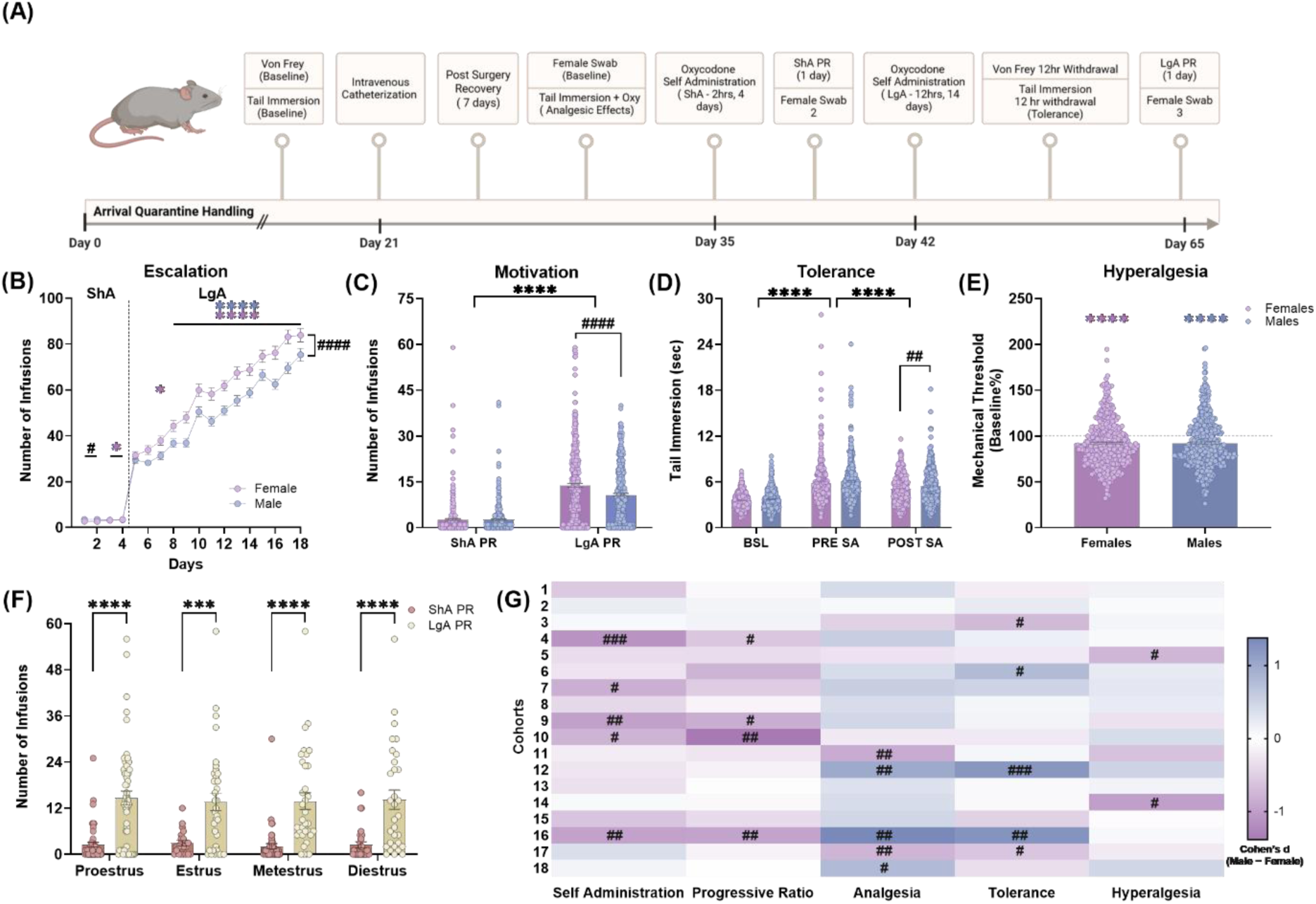
Experimental timeline and sex differences in oxycodone-related behavioral outcomes. **(A)** Experimental timeline for behavioral testing and oxycodone exposure. **(B)** Oxycodone infusions across short - access (ShA; 2 h/day for 4 days) and long-access (LgA; 12 h/day for 14 days) sessions in female (*n* = 384) and male (*n* = 423) HS rats. Colored asterisks indicate significant within-sex increases in intake relative to the first session of the corresponding access condition; # indicates significant between-sex comparisons on days 1,2, 9,11,12,13,14,16, and 17. **(C)** Progressive-ratio (PR) responding following ShA (ShA PR) and LgA (LgA PR) in female (*n* = 387) and male (*n* = 423) HS rats. Asterisks indicate timepoint comparisons; # indicates between-sex comparisons. **(D)** Tail-withdrawal latency at baseline (BSL), following oxycodone administration before self-administration (PRE-SA), and following extended oxycodone self-administration (POST-SA) in female (*n* = 387) and male (*n* = 423) HS rats. Asterisks indicate comparisons across timepoints; # indicates between-sex comparisons. **(E)** Von Frey withdrawal thresholds expressed as percentage of baseline following oxycodone withdrawal in female (*n* = 386) and male (*n* = 423) HS rats. Asterisks indicate comparisons with the theoretical baseline of 100%. **(F)** PR responding across estrous phases following ShA and LgA. ShA PR: *n* = 152 (proestrus, *n* = 55; estrus, *n* = 21; metestrus, *n* = 48; diestrus, *n* = 28); LgA PR: *n* = 154 (proestrus, *n* = 53; estrus, *n* = 36; metestrus, *n* = 34; diestrus, *n* = 31). * Symbols indicate ShA versus LgA comparisons within estrous phase. **(G)** Cohort-level sex differences across behavioral measures. Cohen’s *d* was calculated as male minus female; negative values indicate higher values in females and positive values indicate higher values in males. # Symbols indicate significant sex differences within individual cohorts based on uncorrected two-sided Welch’s t-tests. * or # *p* < 0.05, ** or ## *p* < 0.01, *** or ### *p* < 0.001, **** or #### *p* < 0.0001, unless otherwise indicated.

Besides total lever presses and drug infusions collected from all sessions, lever press data from the final four long-access sessions (LgA days 11–14) was analyzed to characterize the temporal organization of oxycodone intake. Loading-phase intake represented the number of oxycodone infusions earned during the first 10 min of each session, whereas titration-phase intake represented the number of infusions earned during the last 6 hours of each session. A burst was defined as a series of consecutive infusions separated by inter-infusion intervals ≤120 s. Mean rewards per burst was calculated as the average number of infusions within each burst, whereas maximum burst size was defined as the greatest number of infusions earned within a single burst. The inter-infusion interval coefficient of variation was calculated as the standard deviation divided by the mean inter-infusion interval across the final four long-access sessions.

### Progressive ratio testing

Motivation for oxycodone reinforcement was assessed with a progressive ratio (PR) schedule after the ShA and LgA phases. The response requirement increased across successive infusions according to the following schedule: 1, 1, 2, 2, 3, 3, 4, 4, 6, 6, 8, 8, 10, followed by single-response increments from 11 to 50 and 10-response increments thereafter (60, 70, 80, 90, and 100). Sessions ended when an animal failed to complete the next response requirement within 60 min, and the highest successfully completed ratio was recorded as the breakpoint (de Guglielmo et al., 2019; Kallupi et al., 2026).

### Thermal analgesia testing (tail immersion)

Thermal nociception was assessed with the tail immersion test (D’amour & Smith, 1941; Janssen et al., 1963). While gently restrained, 1 cm of the tail was immersed in a 52°C water bath, and withdrawal latency was recorded (de Guglielmo et al., 2014). Testing occurred at three timepoints: baseline (BSL; pre-drug), 15 min after two intravenous oxycodone infusions (150 µg/kg) prior to the start of ShA sessions (Pre**-**SA), and 12 h into withdrawal following 14 LgA sessions (Post-SA) using the same infusion protocol as before. Changes in withdrawal latency from baseline to early post-infusion were used to assess the analgesic effect of oxycodone (Pre-SA – BSL), and reductions in this effect over time were used to assess Tolerance (Pre SA – Post SA).

### Mechanical nociceptive testing (von Frey)

Mechanical sensitivity was measured using a dynamic plantar aesthesiometer (Ugo Basile, Italy) to determine paw withdrawal thresholds (Chaplan et al., 1994; Smith et al., 2020, 2021). Three trials per hind paw were conducted, recording withdrawal latency and force. Testing occurred at baseline (pre-drug) and during acute withdrawal (10–12 h after the final oxycodone session). Percent change from baseline paw withdrawal threshold was calculated to quantify mechanical hyperalgesia.

### Data Analysis

Raw data were evaluated for completeness prior to analysis. Animals that did not complete the behavioral protocol because of catheter failure or death were excluded, following exclusion procedures used in prior studies (de Guglielmo et al., 2024; Sneddon et al., 2026). Animals with otherwise usable but incomplete data were retained in domain-specific analyses when the required data were available. To integrate multiple behavioral domains into a unified measure of addiction variability, an Addiction Index (AI) was calculated for rats with complete data across all behavioral measures. Accordingly, 708 of 810 rats were included in analyses where we assessed the AI (females, N= 338; males, N=370). The AI incorporated four behavioral measures: escalation of oxycodone intake, motivation to obtain oxycodone, tolerance to oxycodone-induced analgesia, and withdrawal-associated hyperalgesia (Belin et al., 2009; Deroche-Gamonet et al., 2004; Kallupi et al., 2020; Sedighim et al., 2021).

Escalation was quantified as the mean number of oxycodone infusions during the final three LgA sessions, and motivation was quantified as the breakpoint obtained during the progressive-ratio test following LgA (LgA PR). Analgesic tolerance was quantified as the decrease in tail-withdrawal latency from the pre-self-administration to the post-self-administration oxycodone test (PRE − POST), with larger values indicating greater tolerance. Withdrawal-associated hyperalgesia was quantified as the percent reduction in mechanical thresholds during withdrawal. For each behavioral domain, Z-scores were computed separately within each cohort and sex, to account for cohort- and sex-specific variability and place all measures on a common scale: 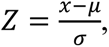 where *x* represents an individual animal’s behavioral value and *μ* and *σ* represent the mean and standard deviation, respectively, for animals of the same sex within that experimental cohort. The AI was calculated as the average value of these four standardized indices, such that higher scores reflect greater addiction-like behavior. AI scores were ranked separately within each sex and divided into quartiles to classify animals into resilient, mild, moderate, and severe groups, representing increasing levels of addiction-like vulnerability.

Behavioral outcomes were analyzed using repeated-measures Analysis of Variance (ANOVA) and mixed-effects models with sex and group as between-subjects factors and repeated measures such as session or timepoint as within-subject factors. Greenhouse-Geisser corrections were applied when the assumption of sphericity was violated. *Post hoc* comparisons were Bonferroni-corrected unless otherwise specified. When data violated parametric assumptions, within-group changes were analyzed using Wilcoxon signed-rank tests and between-group differences with Welch’s t-tests. For comparisons among AI groups, the resilient group served as the reference group where applicable. Estrous-phase effects on PR responding were assessed using a mixed-effects model with estrous phase and timepoint as factors.

Spearman’s rank correlations (ρ) were calculated separately for females and males to examine associations between behavioral measures and addiction-related indices. All correlation coefficients were displayed in the primary correlation heatmaps (Fig. 3), regardless of statistical significance. To identify significant associations within each sex, p-values were adjusted using the Bonferroni correction across 135 behavioral measure–index comparisons separately for females and males. Correlations that remained significant after correction were displayed in Supplementary Figure S1. Differences in correlation strength between sexes were assessed using two-tailed Fisher’s r-to-z tests, with Bonferroni correction applied across the 135 between-sex comparisons. Adjusted p-values were used to identify significant sex differences in correlation strength in both figures.

Within-session oxycodone intake during late long-access self-administration was characterized using loading-phase intake, titration-phase intake, number of bursts, maximum burst size, mean rewards per burst, and inter-infusion interval coefficient of variation (III CV). Sex differences in these intake-pattern measures were evaluated using Welch’s *t*-tests in the full sample, including all rats with available data for each measure. Associations between each intake-pattern measure and composite AI scores were assessed using Spearman’s rank correlations within the AI subset and separately in females and males. Sex differences in correlation strength were evaluated using two-tailed Fisher’s *r*-to-*z* tests. Relationships among intake-pattern measures were also assessed using Spearman’s rank correlations across all available rats and separately by sex, using available paired observations for each comparison. Sex differences in these correlations were evaluated using two-tailed Fisher’s *r*-to-*z* tests. *P*-values were adjusted for multiple comparisons using the Bonferroni correction. To assess the consistency of sex differences across cohorts, cohort-level Cohen’s *d* effect sizes were calculated for each intake-pattern measure using all available rats with data for that measure. Effect sizes were defined in the male-minus-female direction, such that negative values indicate higher values in females. Pooled effects and 95% confidence intervals were estimated using random-effects meta-analysis with restricted maximum likelihood estimation, and heterogeneity was quantified using I².

Cohen’s *d* effect sizes were calculated separately for each behavioral outcome within each cohort. Random-effects meta-analyses were used to estimate pooled effect sizes and 95% confidence intervals across cohorts for each behavioral outcome. Between-cohort heterogeneity was quantified using *I*² and τ². Cohort-level effect sizes and random-effects of pooled estimates were also calculated for the within-session intake-pattern measures.

To assess the magnitude and consistency of sex differences across experimental cohorts, a single value was derived for each behavioral measure for each animal. Self-administration was represented by the mean of 17 overlapping two-session intake bins across the 18 self-administration sessions, progressive-ratio responding by the LgA PR breakpoint, analgesia by the change in tail-withdrawal latency from baseline to the pre-self-administration oxycodone test (PRE − BSL), tolerance by the change from pre- to post-self-administration oxycodone testing (PRE − POST), and hyperalgesia by withdrawal threshold expressed as a percentage of baseline (WD/BSL%). Cohen’s *d* was calculated separately within each cohort in the male-minus-female direction, such that negative values indicate higher measured values in females and positive values indicate higher measured values in males. Cohort-specific effect sizes were combined using random-effects meta-analysis with restricted maximum-likelihood estimation, and between-cohort heterogeneity was quantified using *I*² and τ². Sample sizes required to detect the observed pooled effect sizes were estimated using two-sided independent-samples t-tests, assuming equal group sizes, 80% power, and α = 0.05.

Statistical analyses and data visualization were performed using GraphPad Prism 11 and RStudio with custom R scripts. Major R packages included dplyr, tidyr, ggplot2, patchwork, afex, emmeans, car, metafor, ComplexHeatmap, and irr.

## Results

### Escalation of Oxycodone Self-Administration Exhibits Sex- and Time-Dependent Differences

During the ShA phase, mixed-effects and repeated-measures analyses examined whether oxycodone self-administration escalated differently between sexes. The mixed-effects model revealed no significant main effects of session (*F*(2.60, 2081) = 2.08, *p* = 0.1107) or sex (*F*(1, 803) = 2.41, *p* = 0.121), but a significant sex × session interaction (*F*(3, 2403) = 6.28, *p* = 0.0003). Bonferroni-adjusted *post hoc* tests showed that males earned significantly more rewards than females on the first two days (*p* < 0.05), with no differences thereafter. Within-sex comparisons indicated that females, but not males, significantly increased intake relative to the first session of ShA (day 1) on days 3 (*p* < 0.036) and 4 (*p* < 0.0083; **Fig. 1B**).

During the LgA phase, a mixed-effects analysis identified significant main effects of session (*F*(7.95, 6242) = 223.2, *p* < 0.0001) and sex (*F*(1, 805) = 15.07, *p* = 0.0001), as well as a significant sex × session interaction (*F*(7.95, 6242) = 2.08, *p* = 0.0345). Bonferroni-adjusted *post hoc* tests indicated that females self-administered significantly more oxycodone than males beginning on day 9 (*p* < 0.05), with the exclusion of days 10,15, and 18 (*ps* > 0.3). Within-sex comparisons confirmed significant escalation of intake in both sexes relative to the first session of LgA (day 5), beginning at LgA session 3 (day 7) in females only (*p* = 0.012) and at LgA session 4 (day 8) in both sexes (females: *p* < 0.0001; males: *p* < 0.001; **Fig. 1B**).

### Motivation for Obtaining Oxycodone Increases Following Extended Access and Is Greater in Females

A mixed-effects analysis revealed significant main effects of timepoint (*F*(1, 775) = 601.5, *p* < 0.0001) and sex (*F*(1, 824) = 14.04, *p* = 0.0002), as well as a significant sex × timepoint interaction (*F*(1, 775) = 15.52, *p* < 0.0001). Bonferroni-adjusted *post hoc* comparisons showed that PR breakpoints increased from ShA PR to LgA PR in both sexes (*p* < 0.0001). While no sex difference was observed at ShA PR (*p* >0.05), females achieved significantly higher breakpoints than males at LgA PR (*p* < 0.0001; **Fig. 1C**).

### Estrous phase did not significantly influence progressive-ratio responding following either short or extended access

A mixed-effects analysis examining estrous phase and timepoint (ShA PR vs. LgA PR) revealed no main effect of phase (*F*(3, 298) = 0.056, *p* = 0.983) and no phase × timepoint interaction (*F*(3, 298) = 0.067, *p* = 0.977), but a significant main effect of timepoint (*F*(1, 298) = 94.73, *p* < 0.0001). Bonferroni-adjusted comparisons showed greater PR responding following LgA than ShA within proestrus, estrus, metestrus, and diestrus (all adjusted *p* ≤ 0.0004), whereas no differences were observed among estrous phases at either timepoint (**Fig. 1F**).

### Repeated Oxycodone Exposure Induces Analgesic Tolerance in Both Sexes

A two-way repeated-measures ANOVA revealed significant main effects of timepoint (*F*(1.776, 1435) = 324, *p* < 0.0001) and sex (*F*(1, 808) = 9.468, *p* = 0.0022), with no significant sex × timepoint interaction (*p* = 0.4299). Bonferroni-adjusted *post hoc* comparisons showed that males exhibited higher withdrawal latencies than females at POST-SA (*p* = 0.0213), but not at BSL (*p* = 0.08) or PRE-SA (*p* = 0.16). Within each sex, withdrawal latencies increased from BSL to PRE-SA and decreased from PRE-SA to POST-SA (*all p* < 0.0001), demonstrating initial oxycodone-induced analgesia followed by the development of analgesic tolerance. When evaluating the derived measures, neither oxycodone-induced analgesia (PRE-SA − BSL; *p* = 0.327) nor analgesic tolerance (PRE-SA − POST-SA; *p* = 0.864) differed significantly between females and males (**Fig. 1D**).

### Oxycodone Withdrawal Produces Mechanical Hyperalgesia in Both Sexes

An unpaired Welch’s t-test revealed no significant sex difference in percent change from baseline to withdrawal of the mechanical threshold (t(806.5) = 0.19, *p* = 0.849). Within-sex one-sample Wilcoxon signed-rank tests showed significant decreases from the theoretical baseline of 100% for females (*p* < 0.0001, median change = –10.92%) and for males (*p* < 0.0001, median change = –12.99%; **Fig. 1E**).

### Sex Differences in Oxycodone-Related Behaviors Vary in Magnitude and Consistency Across Cohorts

To rank the magnitude and assess the cross-cohort consistency of sex differences, cohort-specific Cohen’s *d* effect sizes were calculated in the male-minus-female direction and combined using random-effects meta-analysis. The strongest pooled sex differences were observed for self-administration (*d* = −0.37, 95% CI [−0.58, −0.16], *p* = 0.0005) and progressive-ratio responding (*d* = −0.34, 95% CI [−0.51, −0.16], *p* = 0.0002), both reflecting higher values in females. Smaller and nonsignificant pooled effects were observed for analgesia (*d* = 0.25, 95% CI [−0.02, 0.52], *p* = 0.0724), tolerance (*d* = 0.08, 95% CI [−0.17, 0.33], *p* = 0.5381), and hyperalgesia (*d* = −0.01, 95% CI [−0.20, 0.18], *p* = 0.8976). Progressive-ratio responding showed the greatest cross-cohort stability (SD = 0.42, *I*² = 27.2%), followed by hyperalgesia (SD = 0.42, *I*² = 37.1%) and self-administration (SD = 0.44, *I*² = 47.7%), whereas tolerance (SD = 0.56, *I*² = 63.2%) and analgesia (SD = 0.59, *I*² = 68.1%) showed greater between-cohort variability.

At the individual-cohort level, uncorrected Welch’s t-tests identified significant sex differences in five cohorts for escalation, four for progressive-ratio responding, five for analgesia, five for tolerance, and two for hyperalgesia. Notably, significant effects for analgesia and tolerance occurred in both directions, indicating variability in the direction of sex differences across cohorts (**Fig. 1G**).

### AI Severity Does Not Modulate the Effect of Sex on Escalation of Oxycodone Self-Administration

Oxycodone self-administration was first examined across short- and long-access sessions. In females, a Two-Way Repeated-Measures ANOVA during ShA revealed significant main effects of AI group (*F*(3, 334) = 3.89, *p* = 0.0093) and session (*F*(3, 1002) = 6.87, *p* < 0.0001), with no significant interaction (*F*(9, 1002) = 0.64, *p* = 0.767). Pairwise comparisons relative to the resilient group indicated that severe females earned significantly more oxycodone rewards (*p* = 0.016). During LgA, significant main effects of AI group (*F*(3, 334) = 26.70, *p* < 0.0001), session (*F*(13, 4342) = 96.35, *p* < 0.0001), and their interaction (*F*(39, 4342) = 4.65, *p* < 0.0001) were observed. Compared to resilient females, all other groups earned significantly more rewards (severe and moderate: *p* < 0.0001; mild: *p* = 0.0026; **Fig. 2A**).

**Figure 2.**
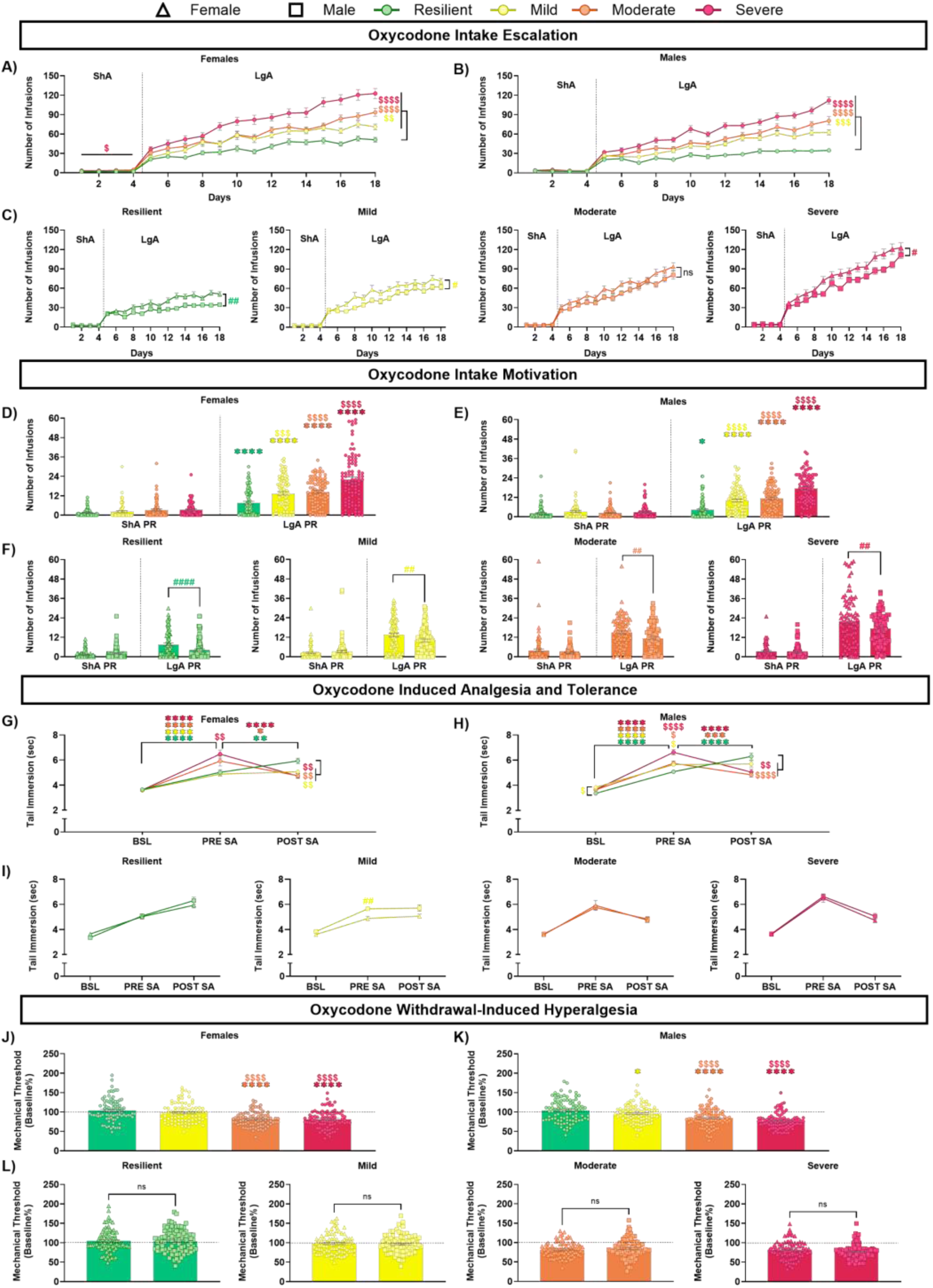
Sex and Addiction-like Severity in Oxycodone-Related Behavioral Outcomes. (Total females: N = 338, Resilient N = 85, Mild N = 84, Moderate N = 84, Severe N =85; Total males: N= 370, Resilient N = 93, Mild N = 92, Moderate N = 92, Severe N = 93). **(A–C)** Number of oxycodone infusions across short access (ShA) and long access (LgA) sessions, shown across AI groups within females (A) and males (B), and between sexes within each AI group (C). **(D–F)** Number of infusions under a progressive-ratio schedule following ShA (ShA PR) and LgA (LgA PR), showing motivation for drug intake across AI groups within females (D) and males (E), and between sexes within each AI group (F). **(G–I)** Oxycodone-induced analgesia and tolerance assessed by tail-withdrawal latency at baseline (BSL), following oxycodone administration before self-administration (PRE-SA), and following extended self-administration (POST-SA), shown across AI groups within females (G) and males (H), and between sexes within each AI group (I). **(J–L)** Oxycodone withdrawal-induced hyperalgesia assessed by von Frey mechanical withdrawal thresholds relative to baseline, shown across AI groups within females (J) and males (K), and between sexes within each AI group (L). Colors indicate AI group (green, resilient; yellow, mild; orange, moderate; red, severe), and symbols indicate sex (triangles, females; squares, males). Data are shown as mean ± SEM with individual animals displayed where applicable. \**p* < 0.05, \*\**p* < 0.01, \*\*\**p* < 0.001, \*\*\*\**p* < 0.0001 indicate timepoint/session differences within groups; $*p* < 0.05, $$*p* < 0.01, $$$*p* < 0.001, $$$$*p* < 0.0001 indicate AI-group differences relative to the resilient group; #*p* < 0.05, ##*p* < 0.01, ###*p* < 0.001, ####*p* < 0.0001 indicate sex differences within AI groups.

In males, a Two-Way Repeated-Measures ANOVA detected no significant effects during ShA (AI group: *F*(3, 366) = 1.68, *p* = 0.170; session: *F*(2.56, 935.9) = 1.59, *p* = 0.198; interaction: *F*(9, 1098) = 1.79, *p* = 0.0659). However, during LgA, significant main effects of AI group (*F*(3, 366) = 41.31, *p* < 0.0001), session (*F*(8.54, 3124) = 114.3, *p* < 0.0001), and their interaction (*F*(39, 4758) = 7.62, *p* < 0.0001) were detected. Pairwise comparisons indicated that all male groups earned significantly more rewards than resilient males (severe and moderate: *p* < 0.0001; mild: *p* = 0.0006; **Fig. 2B**).

To determine whether oxycodone intake differed by sex within each AI group, a two-way ANOVA (sex × session) was performed separately for each group. During ShA, a significant main effect of session was detected in the mild group (*F*(2.374, 413.0) = 3.76, *p* = 0.0108). A significant sex × session interactions were observed in the resilient (*F*(3, 528) = 2.75, *p* = 0.042) and severe (*F*(3, 528) = 2.97, *p* = 0.031) groups; however, Bonferroni-adjusted comparisons revealed no significant sex differences at any individual ShA session (all adjusted *p* > 0.05; **Fig. 2C**).

In contrast, during LgA, main effects of sex were observed in all four AI groups before correction. Following Bonferroni correction across the four AI-group-specific sex effects, females exhibited greater oxycodone self-administration than males in the resilient (*F*(1, 176) = 10.46, adjusted *p* = 0.006), mild (*F*(1, 174) = 7.06, adjusted *p* = 0.034), and severe (*F*(1, 176) = 7.52, adjusted *p* = 0.027) groups, whereas the sex effect in the moderate group did not survive correction (*F*(1, 174) = 5.42, adjusted *p* = 0.084). A significant sex × session interaction was observed only in the resilient group (*F*(13, 2288) = 2.85, *p* = 0.0004), whereas no significant interactions were detected in the other AI groups (**Fig. 2C**). Additional analyses revealed no significant sex × AI group or sex × AI group × session interactions during either ShA (both *p*s ≥ 0.394) or LgA (both *p*s ≥ 0.607) (data not shown).

### AI Severity does not Modulate the Effect of Sex on Progressive Ratio (PR)

In females, a Two-Way Repeated-Measures ANOVA revealed significant main effects of AI group (*F*(3, 334) = 26.5, *p* < 0.0001) and timepoint (*F*(1, 332) = 370.2, *p* < 0.0001), as well as significant AI group × timepoint interactions (*F*(3, 332) = 17.3, *p* < 0.0001). Similarly, males showed significant main effects of AI group (*F*(3, 366) = 36.74, *p* < 0.0001) and timepoint (*F*(1, 366) = 309.1, *p* < 0.0001), along with a significant AI group × timepoint interaction (*F*(3, 366) = 31.04, *p* < 0.0001). No group differences were detected at ShA PR in either sex (all *p* > 0.3). At LgA PR, animals classified as severe, moderate, or mild exhibited significantly elevated breakpoints compared to resilient animals (both sexes: all *p* < 0.0001). Within-sex *post hoc* comparisons confirmed significant increases in breakpoints from ShA PR to LgA PR across all groups in females (all *ps* < 0.0001) and mild, moderate, and severe animals in males (all *p* < 0.0001; **Fig. 2D-E**).

To directly assess sex differences within each AI group, separate two-way ANOVAs (Sex × Timepoint) were performed with Bonferroni-corrected *post hoc* comparisons. No significant sex differences were detected at ShA PR across AI groups. In contrast, females exhibited significantly higher breakpoints than males at LgA PR in all four groups (resilient: *p* < 0.0001; mild: *p* = 0.0046; moderate: *p* = 0.0029; severe: *p* = 0.003). In the resilient group, significant main effects of sex (*F*(1, 176) = 5.95, *p* = 0.0157) and timepoint (*F*(1, 176) = 55.3, *p* < 0.0001), as well as a sex × timepoint interaction (*F*(1, 176) = 12.14, *p* = 0.0006), were observed. In the mild group, there was a significant main effect of timepoint (*F*(1, 174) = 125.9, *p* < 0.0001) and a sex × timepoint interaction (*F*(1, 174) = 7.33, *p* = 0.008), but no main effect of sex. In the moderate group, significant main effects of sex (*F*(1, 174) = 9.56, *p* = 0.0023) and timepoint (*F*(1, 174) = 156.8, *p* < 0.0001) were observed without a significant interaction. Finally, in the severe group, significant main effects of sex (*F*(1, 176) = 5.96, *p* = 0.0162) and timepoint (*F*(1, 176) = 314.2, *p* < 0.0001), as well as a sex × timepoint interaction (*F*(1, 176) = 4.29, *p* = 0.04), were observed (**Fig. 2F**). Additional analyses revealed no significant sex × AI group or sex × AI group × timepoint interactions (both *p*s ≥ 0.524).

### AI Severity does not Modulate the Effect of Sex on Tolerance to Oxycodone Analgesia

Tolerance to oxycodone analgesia was examined across AI groups. Separate two-way repeated-measures ANOVAs revealed significant main effects of timepoint in both females (*F*(1.656, 553.1) = 118.5, *p* < 0.0001) and males (*F*(1.939, 709.8) = 209.4, *p* < 0.0001), as well as significant timepoint × AI group interactions (females: *F*(6, 668) = 11.22, *p* < 0.0001; males: *F*(6, 732) = 15.61, *p* < 0.0001). No significant main effect of AI group was observed in either females (*F*(3, 334) = 1.69, *p* = 0.168) or males (*F*(3, 366) = 1.53, *p* = 0.206).

At BSL, no significant differences between AI groups were detected in females, whereas mild males exhibited higher withdrawal latencies than resilient males (*p* = 0.015). At PRE-SA, mild (*p* = 0.039), moderate (*p* = 0.025), and severe (*p* < 0.0001) males exhibited higher withdrawal latencies than resilient males. In females, severe animals exhibited higher PRE-SA withdrawal latencies than resilient animals (*p* =0.0007). At POST-SA, resilient animals exhibited higher withdrawal latencies than moderate (females: *p* = 0.0002; males: *p* < 0.0001) and severe animals (females: *p* = 0.0001; males: *p* = 0.0022). Resilient females also exhibited higher withdrawal latencies than mild females (*p* = 0.005), whereas resilient and mild males did not significantly differ.

Within each sex, withdrawal latencies increased significantly from BSL to PRE-SA across all AI groups (all *ps* < 0.0001), indicating oxycodone-induced analgesia. The subsequent change from PRE-SA to POST-SA differed across vulnerability groups. In both sexes, withdrawal latencies increased further in the resilient group (females: *p* = 0.0025; males: *p* < 0.0001), did not significantly change in the mild group (both *p* > 0.9999), and decreased in the moderate (females: *p* = 0.0128; males: *p* = 0.0002) and severe groups (both *p* <0.0001), consistent with greater analgesic tolerance in the moderate and severe groups. To directly assess sex differences within each AI group, separate two-way repeated-measures ANOVAs (sex × timepoint) were performed. Significant main effects of timepoint were observed in all groups (all *ps* < 0.0001), whereas no significant sex × timepoint interactions were detected (all *ps* > 0.15). Significant main effect of sex was only observed in the mild group (*F*(1, 174) = 9.59, *p* = 0.0023), with males exhibiting higher withdrawal latencies overall. Bonferroni-corrected comparisons indicated higher withdrawal latencies in males than females at PRE-SA (*p* = 0.0035), but not at BSL (*p* = 0.539) or POST-SA (*p* = 0.110). Further three way Anova revealed no significant sex × AI group or sex × AI group × timepoint interactions (both *p*s ≥ 0.179; **Fig. 2G-I**).

### AI Severity does not Modulate the Effect of Sex on Withdrawal-Induced Mechanical Hyperalgesia

In females, Bonferroni-adjusted Welch’s *t*-tests comparing each AI group with the resilient group showed that moderate and severe groups exhibited significantly greater hyperalgesia (both *ps* < 0.0001), whereas the mild and resilient groups did not differ (*p* = 0.594). Within-group Wilcoxon signed-rank tests against the theoretical baseline (100 percent) confirmed significant hyperalgesia in the moderate and severe groups (both *p* < 0.0001), but not in the mild (*p* = 0.223) or resilient (*p* = 0.339) groups. In males, group differences followed a similar pattern. Moderate and severe males exhibited greater hyperalgesia compared to resilient males (both *p* < 0.0001). Within-group Wilcoxon tests indicated significant hyperalgesia in mild (*p* = 0.0374), moderate (*p* < 0.0001), and severe (*p* < 0.0001) males, but not in the resilient group (*p* = 0.266; **Fig. 2J-K**). Sex comparisons using Welch’s *t*-tests within each AI group revealed no statistically significant differences: resilient (*p* = 0.909), mild (*p* = 0.586), moderate (*p* = 0.315), and severe (*p* = 0.115; **Fig. 2L**). Consistent with these comparisons, there was no significant sex × AI group interaction (*p* = 0.440).

### Relationships Among Addiction-Related Behaviors Are Largely Conserved Across Sexes

To characterize relationships among addiction-related behavioral measures, Spearman correlation analyses were performed separately in females (n = 338) and males (n = 370), combining animals across all AI groups (**Fig. 3**). Overall, correlation patterns were largely similar between sexes, and strong correlations were observed between individual behavioral measures and their corresponding indices. Beyond these expected associations, average oxycodone intake during the final extended-access sessions was significantly correlated with the Motivation Index in females (ρ = 0.455) and males (ρ = 0.477), whereas correlations with the Tolerance and Pain Indices were generally weak in both sexes.

**Figure 3.**
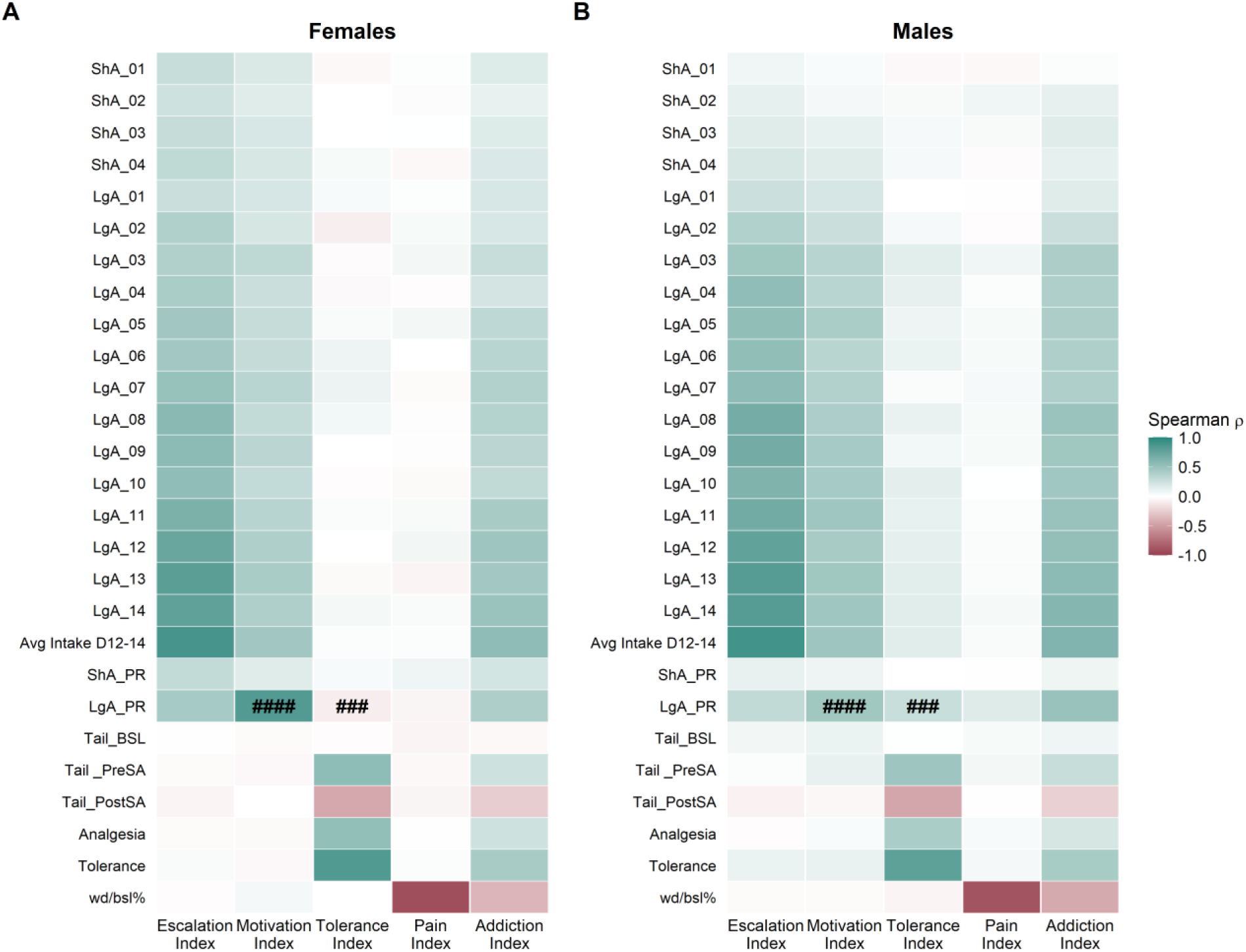
Sex-specific correlations between oxycodone-related behavioral measures and addiction-related indices. Spearman correlation heatmaps showing associations between behavioral measures and the Escalation, Motivation, Tolerance, Pain, and Addiction Indices in (A) females (n = 338) and (B) males (n = 370), with all AI groups combined. Colors represent the direction and strength of Spearman’s correlation coefficients (ρ). All correlations are displayed regardless of statistical significance. Differences in correlation strength between sexes were assessed using Fisher r-to-z approximations with Bonferroni correction. Symbols indicate significant sex differences in correlation strength (#p < 0.05, ##p < 0.01, ###p < 0.001, ####p < 0.0001; Bonferroni-adjusted). Symbols appear in both panels for corresponding correlations.

Two sex differences in correlation strength remained significant after Bonferroni correction. The association between LgA PR and the Motivation Index was stronger in females (ρ = 0.838) than males (ρ = 0.461; adjusted p < 0.0001). In contrast, LgA PR was positively associated with the Tolerance Index in males (ρ = 0.273), whereas a weak negative association was observed in females (ρ = −0.088; adjusted p = 0.00015). No other sex differences in correlation strength survived correction for multiple comparisons (**Fig. 3**). Supplementary Figure S1 presents the same correlation analyses, displaying only associations that remained significant after Bonferroni correction.

### Oxycodone Intake Patterns Vary by Sex and Addiction Variability

Having characterized sex- and addiction variability-related differences across the primary addiction-like behavioral domains, we next examined the temporal organization of oxycodone intake during late LgA sessions 11–14 (days 15–18). Sex differences in these measures were assessed using Welch’s unpaired *t*-tests, with *p*-values adjusted across the six measures using the Bonferroni correction. Females exhibited a greater number of bursts (*t*(792.4) = 4.18, adjusted *p* = 0.0002) and greater maximum burst size (*t*(649.4) = 3.29, adjusted *p* = 0.0062) than males. Sex differences in loading-phase intake, titration-phase intake, mean rewards per burst, and inter-infusion interval coefficient of variation were not significant (**Fig. 4A–F**).

**Figure 4.**
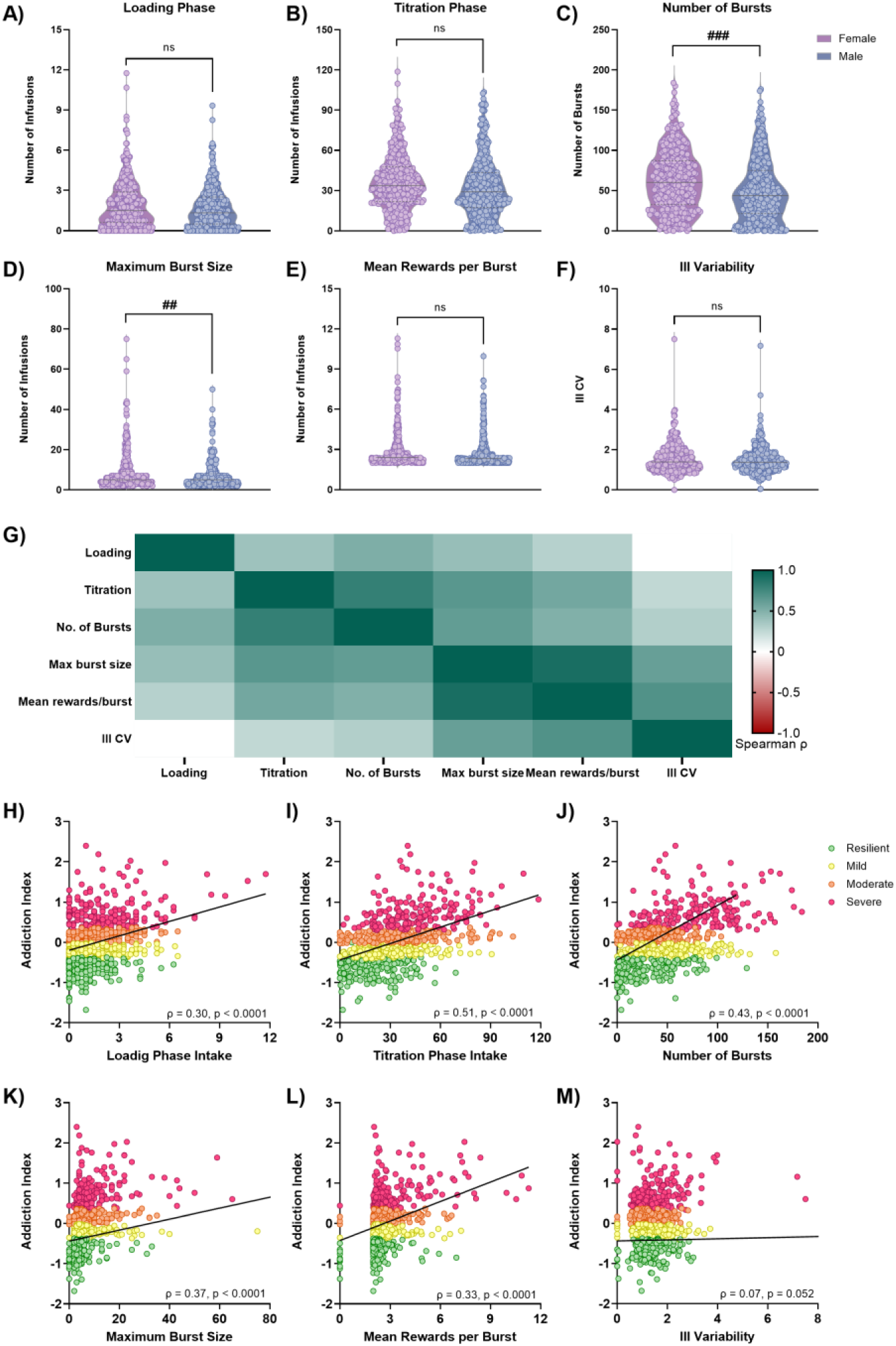
Sex differences and relationships with addiction-like severity in temporal patterns of oxycodone intake. **(A–F)** Sex differences in within-session oxycodone intake patterns calculated across the final four long-access sessions (LgA days 11–14), including loading-phase intake **(A)**, titration-phase intake **(B)**, number of bursts **(C)**, maximum burst size **(D)**, mean rewards per burst **(E)**, and inter-infusion interval coefficient of variation (III CV) **(F)**. Sample sizes varied according to data availability: loading and titration, *n* = 798; number of bursts, *n* = 803; maximum burst size and mean rewards per burst, *n* = 786; III CV, *n* = 788. Sex differences were assessed using Welch’s *t*-tests with Bonferroni correction across measures. **(G)** Spearman correlation matrix of the six temporal intake measures. Colored cells indicate correlations that survived Bonferroni correction, whereas white cells indicate correlations that did not survive correction. **(H–M)** Spearman correlations between the Addiction Index (AI) and loading **(H)**, titration **(I)**, number of bursts **(J)**, maximum burst size **(K)**, mean rewards per burst **(L)**, and III CV **(M)** (*n* = 704). Points are colored according to AI group (resilient, mild, moderate, and severe). # sign indicates significant sex differences following Bonferroni correction (# *p* < 0.05, ## *p* < 0.01, ###*p* < 0.001).

To assess the consistency of these sex differences across cohorts, cohort-level effect sizes were examined using random-effects meta-analysis. Pooled effects were consistent with the overall sex comparisons, with significant female-biased effects for number of bursts and maximum burst size, whereas effects for loading-phase intake, titration-phase intake, mean rewards per burst, and III CV were not significant after Bonferroni correction. Heterogeneity across cohorts ranged from *I*² = 0% to 50.2% (**Fig. S2**).

To determine how the different features of oxycodone intake were related to one another, we next examined correlations among the intake-pattern measures. Intake-pattern measures were broadly positively correlated with one another during late LgA. The strongest association was observed between maximum burst size and mean rewards per burst (ρ = 0.93), followed by titration-phase intake and number of bursts (ρ = 0.84), titration-phase intake and maximum burst size (ρ = 0.65), and number of bursts and maximum burst size (ρ = 0.64; all adjusted *ps* < 0.0001). All other pairwise associations were also significant after Bonferroni correction (all adjusted *ps*< 0.0001), except for the relationship between loading-phase intake and III CV (ρ = 0.04, adjusted *p*= 1; **Fig. 4G**). Sex-stratified analyses revealed broadly similar relationships among intake-pattern measures in females and males. The strongest association in both sexes was between maximum burst size and mean rewards per burst (females: ρ = 0.91; males: ρ = 0.93), followed by titration-phase intake and number of bursts (females: ρ = 0.80; males: ρ = 0.86; all adjusted *ps* < 0.0001). All other pairwise associations were also significant after Bonferroni correction, except for the relationship between loading-phase intake and III CV (females: ρ = 0.06, adjusted *p* = 1; males: ρ = 0.02, adjusted *p* = 1). Direct comparisons of correlation strength revealed no significant sex differences after Bonferroni correction across the 15 pairwise comparisons (all adjusted *p* ≥ 0.091). The association between titration-phase intake and number of bursts was nominally stronger in males than females (*z* = −2.73, *p* = 0.006), but this difference did not survive Bonferroni correction (adjusted *p* = 0.091).

Having characterized individual variation in the organization of oxycodone intake during late LgA, we next examined whether these intake patterns were associated with AI scores. Across all animals, Spearman’s rank correlations revealed that titration-phase intake showed the strongest positive association with AI scores (ρ = 0.51, *p* < 0.0001), followed by number of bursts (ρ = 0.49, *p* < 0.0001), maximum burst size (ρ = 0.37, *p* < 0.0001), mean rewards per burst (ρ = 0.33, *p* < 0.0001), and loading-phase intake (ρ = 0.30, all Bonferroni-adjusted *p* < 0.0001). In contrast, III CV was not significantly associated with AI scores (ρ = 0.07, adjusted *p* = 0.315; **Fig. 4 H-M**). These associations were broadly similar between females and males, with no significant sex differences in correlation strength after Bonferroni correction (all adjusted *p* ≥ 0.160; **Fig. S3**).

## Discussion

The present study examined how sex and individual variability contribute to multiple dimensions of oxycodone addiction-like behaviors in a genetically diverse rat population. First, sex differences in oxycodone self-administration and PR responding were dependent on the stage of drug exposure, with females exhibiting greater oxycodone intake and motivation following extended access. Second, self-administration and PR responding were the behavioral measures most sensitive to sex differences, with PR responding showing the greatest consistency across cohorts, whereas analgesic tolerance and withdrawal-induced hyperalgesia exhibited comparatively limited sex effects. However, additional analyses provided no evidence that the magnitude of sex differences varied across AI groups. Finally, analyses revealed greater number of bursts and larger maximum burst sizes during extended access, whereas correlations among addiction related behaviors and within-session intake patterns were largely similar between sexes.

Sex differences in oxycodone self-administration varied across the course of drug exposure. During ShA, males initially exhibited greater intake than females, whereas females increased their intake across sessions. Following transition to LgA, intake progressively increased in both sexes, with females exhibiting greater intake than males later in extended access. These findings are consistent with previous studies demonstrating greater opioid self-administration and escalation in females under extended-access conditions. Importantly, the present findings extend this literature by demonstrating that sex differences in oxycodone intake depend on the stage of drug exposure and occur across individual addiction-like phenotypes. Although sex × session interactions were observed within selected AI groups, the absence of significant sex × AI group and sex × AI group × session interactions indicate that the magnitude and temporal pattern of sex differences did not significantly vary across AI groups. Thus, individual variability in addiction-like behavior was strongly associated with oxycodone intake but did not significantly alter the overall expression of sex differences in self-administration.

A similar exposure-dependent pattern was observed for progressive-ratio responding. Progressive-ratio responding increased following extended access in both sexes, while sex differences were absent following short access and emerged following extended access, when females exhibited greater responding than males. Females also exhibited greater progressive-ratio responding following extended access within each addiction index group. This increased female responding is consistent with previous studies reporting greater opioid progressive-ratio responding in females (Becker & Hu, 2008; Kallupi et al., 2026; Mavrikaki et al., 2017). However, the absence of significant sex × AI group and sex × AI group × timepoint interactions indicate that these sex differences did not significantly vary in magnitude or across timepoints among addiction index groups. Together, these findings indicate that extended oxycodone exposure is an important context for the emergence of sex differences in progressive-ratio responding, whereas individual addiction-like variability was associated with the overall level of responding in both sexes rather than differential expression of the sex effect across AI groups.

Estrous phase did not significantly influence PR responding following either short or extended access, nor did estrous phase alter the increase in PR responding observed following extended access. This finding adds to a mixed literature in which estrous-dependent differences in opioid self-administration have varied across drug, dose, and access conditions (Cicero et al., 2003; Hinds et al., 2023; Lynch & Carroll, 1999; Patel & Loweth, 2024; Sharp et al., 2021; Towers et al., 2023). One factor that may contribute to the lack of an association in the present study is the substantial variability in estrous cyclicity observed in HS rats. Sneddon et al. (2026) reported frequent irregular or prolonged estrous phases in this population, suggesting that cytologically defined phase may not consistently correspond to stereotypical hormonal profiles. The absence of a phase effect in the present study does not establish that ovarian hormones are unrelated to opioid-related behavior but rather indicates that estrous phase at the time of PR testing was not associated with PR responding under these conditions.

Pain-related behavioral outcomes showed a different relationship with individual addiction-like variability. Oxycodone produced robust analgesia in both sexes, followed by reduced analgesic responsiveness after extended self-administration, consistent with the development of tolerance and consistent with our previous work (Kallupi et al., 2026). The trajectory of this response differed across AI groups: resilient animals exhibited increased analgesic responsiveness following extended access, whereas only moderate and severe animals showed reduced analgesic responsiveness, consistent with the development of tolerance. Withdrawal-induced mechanical hyperalgesia was similarly differentiated by AI groups, with greater hyperalgesia in the moderate and severe groups. These findings demonstrate that individual differences captured by the AI extend beyond drug-taking behavior and are also reflected in adaptations related to opioid analgesia and withdrawal-associated pain.

Sex differences in these pain-related measures were comparatively limited. Although sex contributed to withdrawal latency in the overall analysis and a specific sex difference was detected at PRE-SA within the mild AI group, the broader temporal pattern of tolerance was similar between females and males. Importantly, the significant PRE-SA difference within the mild group does not indicate that the sex effect was uniquely greater in this AI group: neither the sex × AI group nor sex × AI group × timepoint interactions were significant. Consistent with our previous work, withdrawal-induced hyperalgesia likewise did not differ significantly between sexes overall (Kallupi et al., 2026; Kimbrough et al., 2020) or within individual AI groups, and there was no sex × AI group interaction. Thus, compared with the increased intake in female with extended-access self-administration and PR responding, pain-related adaptations were more strongly differentiated across AI groups than between sexes.

The cohort-level analyses further characterized the magnitude and consistency of sex differences across the 18 experimental cohorts. Pooled effect sizes indicated significantly greater self-administration and PR responding in females, consistent with the overall behavioral analyses. These were the strongest pooled sex effects, with PR responding exhibiting the lowest between-cohort heterogeneity, suggesting that sex differences in these behaviors were relatively consistent across cohorts. Based on the observed pooled effect sizes, power analyses estimated that approximately 115 animals per sex would be required to detect sex differences in self-administration and 141 per sex for PR responding, assuming 80% power and a two-sided α of 0.05. In contrast, pooled sex effects for analgesia, tolerance, and hyperalgesia were not significant, and their smaller effect sizes yielded substantially larger sample-size estimates. Greater between-cohort heterogeneity was observed for analgesia and tolerance, with significant sex differences occurring in both directions across individual cohorts.

To our knowledge this is the first study to investigate individual- and sex-difference across multiple cohorts that comprehensively analyzes these oxycodone-related behavioral profile (outside of our previous work, Kallupi et al., (2026) which this work extends on). These findings complement the overall behavioral comparisons by highlighting relatively consistent female-biased effects in drug taking and motivation, whereas sex differences in pain-related measures were more variable and less consistently associated with one sex. Relationships among the primary behavioral measures and addiction-related indices further highlighted the multidimensional organization of addiction-like behavior. Individual behaviors were strongly associated with their corresponding indices and with the composite AI, consistent with their contribution to index construction. Importantly, cross-domain relationships were also observed, including associations between late-LgA intake and the Motivation Index and between LgA PR and the Escalation Index. These findings suggest that escalation and PR responding capture related, but not identical, aspects of oxycodone self-administration, consistent with addiction-like behavior reflecting coordinated variation across multiple behavioral domains rather than a single behavioral measure.

Although the overall correlation structure was broadly similar between females and males, two specific relationships differed by sex. Notably, the association between LgA PR and the Motivation Index was stronger in females, whereas the relationship between LgA PR and the Tolerance Index was positive in males but not females. Together, these findings suggest that sex is associated with selective differences in relationships among addiction-related behaviors rather than widespread differences in the overall correlation patterns.

Beyond the primary behavioral domains, the temporal organization of oxycodone intake provided additional information about sex differences during extended access. Females exhibited greater number of bursts, and maximum burst size during the final LgA sessions, whereas loading-phase intake, titration-phase intake, mean rewards per burst, and III coefficient of variation did not significantly differ between sexes after correction. These findings are consistent with recent work showing sex-dependent differences in burst-like oxycodone intake during escalation (Hodges et al., 2026) and with prior studies highlighting front-loading as a behaviorally relevant feature of oxycodone self-administration (Guha et al., 2022). Together, these studies highlight the value of examining within-session intake structure in addition to total drug consumption.

Cohort-level analyses of these measures showed generally female-biased effects, providing additional evidence that sex may influence not only cumulative oxycodone intake but also the underlying behavioral processes reflected in the temporal distribution of drug consumption.

Several temporal intake measures were also associated with the composite AI, with titration-phase intake showing the strongest relationship, followed by the number of bursts and maximum burst size. Titration and burst measures were strongly related, indicating that sustained late-session intake was associated with greater clustering of infusions. In contrast, III coefficient of variation was not significantly associated with AI in the overall population. Because the within-session intake measures and the escalation component of the AI were derived from late extended-access sessions, their correlations may partly reflect shared variation in oxycodone consumption.

Our findings demonstrate that animals with higher AI scores also differed in how oxycodone intake was distributed within extended-access sessions. Notably, direct comparisons of the relationships among temporal intake measures revealed few sex differences after correction for multiple comparisons. Thus, sex differences were more apparent in the magnitude of specific intake patterns than in the relationships among those patterns, suggesting that the broader organization of within-session oxycodone self-administration was largely shared across sexes.

Some limitations should be considered. Estrous phase was determined using vaginal cytology rather than direct measurement of circulating ovarian hormones, limiting conclusions regarding specific hormonal contributions to PR responding. Tail immersion and von Frey testing assess thermal analgesia and mechanical sensitivity, respectively, and therefore do not encompass the multidimensional features of pain associated with chronic opioid exposure. In addition, several behavioral measures were used in the construction of the AI or its component indices; consequently, associations between these measures and the AI partly reflect shared mathematical and behavioral information and should not be interpreted as independent prediction.

Overall, sex and individual variability contributed in distinct and behavior-dependent ways to oxycodone addiction-like behavior. Female-biased behavioral differences were most apparent following extended access, including greater oxycodone intake, PR responding, number of bursts, and maximum burst size. In contrast, tolerance and withdrawal-induced hyperalgesia were more strongly differentiated across AI groups than between sexes. Importantly, although specific sex differences were observed within each AI group, further analyses provided no evidence that their magnitude varied across AI groups. Behavioral relationships were also largely shared between sexes, with selective differences in specific associations rather than broad reorganization of the behavioral structure. Together, these findings provide a multidimensional framework for understanding how sex and individual variability contribute to opioid addiction-like behavior in a genetically diverse population.

