## Supplementary Material for "Severity of Addiction-like Behaviors and Sex Distinctly Promote Oxycodone-Related Behaviors in Heterogeneous Stock Rats"

Foroutani et al.

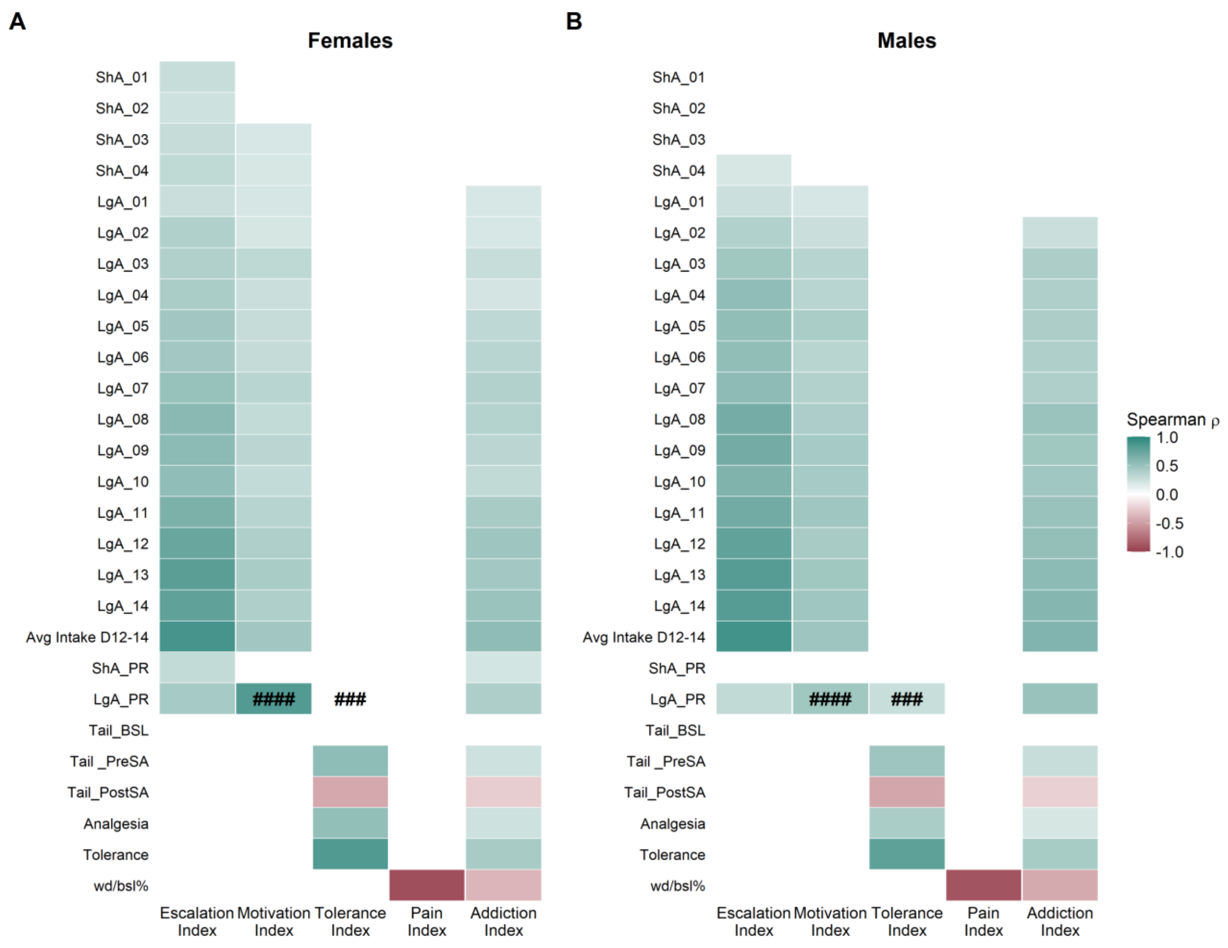


**Figure S1. Bonferroni-significant correlations between oxycodone-related behavioral measures and addiction-related indices by sex.** Spearman correlation heatmaps showing associations between behavioral measures and the Escalation, Motivation, Tolerance, Pain, and Addiction Indices in (A) females (n = 338) and (B) males (n = 370), with all Addiction Index (AI) groups combined. Only correlations that remained significant after Bonferroni correction for 135 behavior–index comparisons within each sex (adjusted $p < 0.05$) are colored. Burgundy represents negative correlations, teal represents positive correlations, and white indicates correlations that did not survive correction or could not be calculated. Symbols indicate significant differences in correlation strength between females and males following Bonferroni correction of Fisher $r$-to-$z$ comparisons across 135 tests (# adjusted $p < 0.05$; ## adjusted $p < 0.01$; ### adjusted $p < 0.001$; #### adjusted $p < 0.0001$). Hash symbols appear in both panels for the corresponding associations.

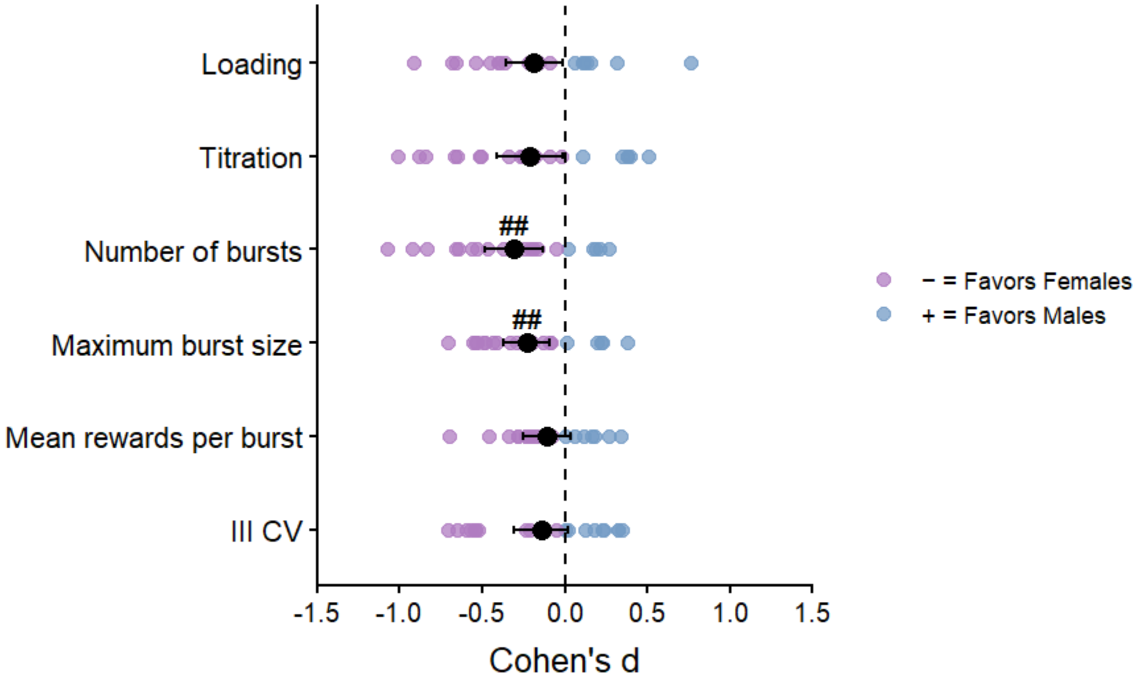


**Figure S2. Cohort-level effect sizes for sex differences in late long-access oxycodone intake patterns.** Negative effect sizes indicate higher values in females and positive effect sizes indicate higher values in males. Colored points represent individual cohort effect sizes; black points and error bars represent pooled estimates and 95% confidence intervals.

Cohort-level Cohen's *d* values were calculated for each intake-pattern measure and combined using random-effects meta-analysis. Following Bonferroni correction across the six intake-pattern measures, pooled effect sizes were significant for number of bursts ($d = -0.31$, 95% CI [−0.48, −0.13], adjusted $p = 0.003$) and maximum burst size ($d = -0.23$, 95% CI [−0.37, −0.09], adjusted $p = 0.008$). Pooled effects were not significant for loading-phase intake, titration-phase intake, mean rewards per burst, or III CV. Heterogeneity ranged from $I^2 = 0\%$ to 50.2%.

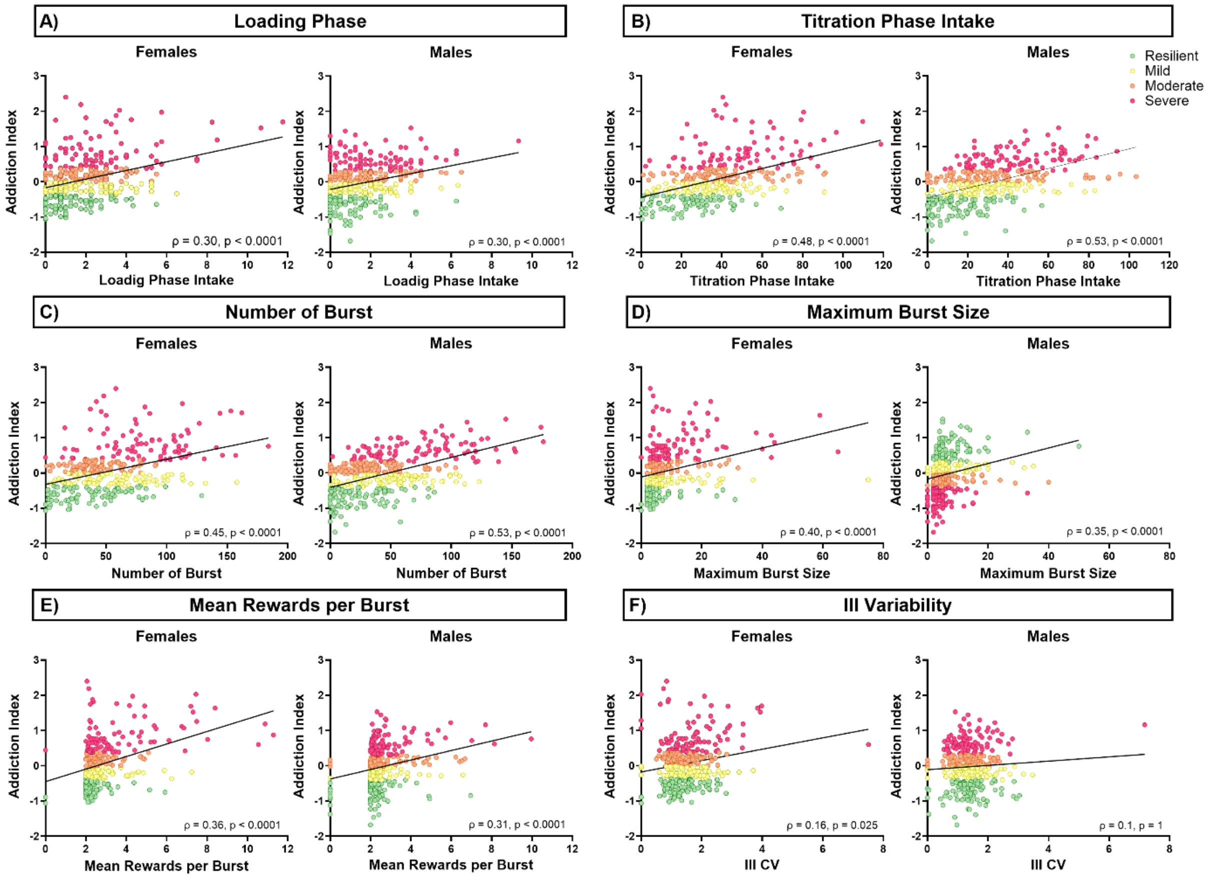


**Figure S3. Sex-specific associations between oxycodone intake patterns and addiction-like vulnerability**

Sex-stratified Spearman's rank correlations showed that (A) loading-phase intake, (B) titration-phase intake, (C) number of bursts, (D) maximum burst size, and (E) mean rewards per burst were positively associated with AI scores in both females and males (all Bonferroni-adjusted *p*s < 0.0001). (F) III CV was positively associated with AI scores in females ($\rho = 0.16$, Bonferroni-adjusted $p = 0.025$), but not in males ($\rho = 0.01$, Bonferroni-adjusted $p = 1.000$). However, direct comparisons of correlation coefficients using Fisher's $r$-to-$z$ tests revealed no significant sex differences following Bonferroni correction across the six intake-pattern measures (all adjusted *p*s $\geq 0.160$). The female–male difference in the association between III CV and AI was significant before correction ($z = 2.22$, $p = 0.027$) but did not remain significant following Bonferroni correction (adjusted $p = 0.160$).
